# Streamlining Computational Fragment-Based Drug Discovery through Evolutionary Optimization Informed by Ligand-Based Virtual Prescreening

**DOI:** 10.1101/2023.11.27.568919

**Authors:** Rohan Chandraghatgi, Hai-Feng Ji, Gail L. Rosen, Bahrad A. Sokhansanj

## Abstract

Recent advances in computational methods provide the promise of dramatically accelerating drug discovery. While mathematical modeling and machine learning have become vital in predicting drug-target interactions and properties, there is untapped potential in computational drug discovery due to the vast and complex chemical space. This paper advances a novel computational fragment-based drug discovery (FBDD) method called Fragment Databases from Screened Ligands Drug Discovery (FDSL-DD), which aims to streamline drug design by applying a two-stage optimization process. In this approach, ***in silico*** screening identifies ligands from a vast library, which are then fragmentized while attaching specific attributes based on predicted binding affinity and interaction with the target sub-domain. This process both shrinks the search space and focuses on promising regions within it. The first optimization stage assembles these fragments into larger compounds using evolutionary strategies, and the second stage iteratively refines resulting compounds for enhanced bioactivity. The methodology is validated across three diverse protein targets involved in human solid cancers, bacterial antimicrobial resistance, and SARS-CoV-2 viral entry, demonstrating the approach’s broad applicability. Using the proposed FDSL-DD and two-stage optimization approach yields high-affinity ligand candidates more efficiently than other state-of-the-art computational methods. Furthermore, a multiobjective optimization method is presented that accounts for druglikeness while still producing potential candidate ligands with high binding affinity. Overall, the results demonstrate that integrating detailed chemical information with a constrained search framework can markedly optimize the initial drug discovery process, offering a more precise and efficient route to developing new therapeutics.

## INTRODUCTION

Advances in computational methodologies have revolutionized drug discovery. Mathematical modeling and machine learning techniques have emerged as vital tools to predict drug-target interactions and drug properties. ^14,30,58,66,69,72,77^ However, computation has yet to be employed to its full potential in the drug discovery process. One key area for innovation is computational drug discovery. ^34,67,85^ As the low-hanging fruit for novel drugs has been harvested, it is increasingly difficult to find promising lead compounds. ^18,19,32^ The now-conventional approach to drug design relies on high-throughput screening of large compound libraries, which is often time consuming and resource intensive. ^50,52,60,80^ Computational drug discovery has the potential to leverage the power of *in silico* screening by using machine learning and optimization techniques to predict the bioactivity of potential com-ounds and, thereby, accelerate drug discovery.

Recently, “fragment-based drug discovery” (or “design”) (FBDD) has emerged as a potentially promising approach. ^10,22,37,40,64,65^ Unlike other drug design strategies (i.e., structure-based drug discovery, SBDD, or ligand-based drug discovery, LBDD), which usually involve designing or testing full-sized drug molecules, works by identifying smaller chemical fragments that bind effectively to target biomolecules. These small fragments serve as building blocks, which can be grown, linked, or merged to create new drug molecules. FBDD offers a flexible and efficient way to explore the vast potential space of drug-like molecules. However, despite the potential advantages of FBDD, a significant challenge remains: the scale of the chemical space to be explored. Given the enormous diversity of possible chemical fragments and the ways they can be combined, the number of potential drug candidates is effectively infinite. This massive combinatorial problem can become a stumbling block, slowing down the drug discovery process and making it difficult to identify promising candidates. This paper proposes a solution to this challenge, centered around computational generation of potential drug molecules.

Computational *de novo* drug design involves the use of techniques such as genetic algorithms ^3,39,47,51,54,73^, reinforcement learning, including deep reinforcement learning ^57,61,63,74,86^, generative deep learning models ^5,6,9,35,41,43,55^, or other deep learning methods, e.g., graph transformers ^46,71^, models that blend deep learning and evolutionary algorithms ^1,26,54^, and string-based transformers (i.e. operating on a SMILES string representation of molecules) ^27,33^. The algorithms “computationally synthesize” novel drug molecules, either by starting from scratch and adding atoms to form a novel molecule or modifying or adding atoms on an existing chemical structure (“scaffold”). The result is the creation of novel molecules by a) simulating chemical modifications that optimize for the single objective of improving binding efficiency to a target or b) multiobjective optimization including druglikeness objectives, e.g., solubility and other drug-likeness factors. ^11,17,24,25,38,42,47,53,84^

In computational FBDD, various *in silico* computational techniques are utilized to construct fragment libraries for Fragment-Based Drug Discovery (FBDD). The conventional approach to computational FBDD involves either computationally fragmentizing a compound (ligand) library or self-generating fragments using computational techniques, followed by computationally docking target fragments to a protein binding pocket and computationally “growing” or synthesizing a candidate ligandby modifying the frag-ment within that pocket. ^7^ Methods like FastGrow emphasize identifying fragment growth points rather than the specifics of fragment expansion, often comparing to other structural docking tools ^59^. In the realm of docking, ultra-large scale docking techniques, such as those by Lyu et al., identify potential molecules based on docking scores, with a breadth possibly surpassing human intuition ^49^. On a similar note, Allen et al. present iterative fragment growth relying on docking scores, yet distinctively using prescreening information to navigate their search ^4^. The advent of “deep evolutionary learning” for FBDD introduced the employment of a latent space grounded in SMILES, as seen in methods like FragVAE, which incorporates evolutionary operators and data augmentation in the process ^26^. Subsequent techniques, such as Podda et al.’s encoder-decoder generative model, also employ the SMILES structure to produce fragments ^62^. More advanced strategies integrate graph-based and evolutionary operators on a molecule’s latent representation, focusing on multi-objective optimization ^53^. While some approaches start from a template and deploy generative methods for modification based on Structure-Activity Relationship (SAR) ^83^, others like Cortes-Cabrera et al. use a fragment approach, progressively constructing a molecule and utilizing the ligand efficiency index (LEI) for guidance ^12^. A notable method introduced by Kerstjens et al. applies new genetic operators to fragment- and graph-based evolutionary designs, emphasizing atom compatibility rules ^36^. Many of these methods, including those that utilize reaction rules or grow rules, emphasize the growth of fragments within 3D binding pockets, a feature that resembles this research’s approach ^48^. Advanced techniques, such as those by Tang et al., combine Deep Reinforcement Learning with chemistry to sculpt fragment libraries ^75^. In the wake of these innovations, researchers are also capitalizing on SMILES using transformers to decorate scaffolds, iterating on their multi-objective techniques as seen in the advancements from Liu et al. ^45,46^.

Despite the sophistication of current algorithms, the vastness of the exploration space and the plethora of potential optima remain daunting challenges. Leveraging either heuristic (evolutionary) approaches or learning techniques, these methods aim to identify superior optima. However, given the expansive nature of the chemical space, any strategy will be fail to be a universally optimal solution to all possible problems and chemical configurations. ^82^ In addition to the limits on heuristic optimization methods, deep learning methods are limited because training typically probes only a minute subspace of the chemical structure landscape. The pivotal question that emerges is: Can we harness problem-specific chemical information to effectively curtail the massively complex search space of potential chemical structures?

To reduce the combinatorial search space and achieve more targeted drug designs, we leverage a pipeline for computational FBDD recently developed by our group called Fragments from Ligands Drug Discovery (FDSL-DD). ^81^ The FDSL pipeline involves an initial *in silico* screening step of a large ligand database against a protein target, utilizing computational docking software, e.g., Autodock VINA. The ligands are then computationally fragmentized, and the fragments are assigned information (i.e., fragment attributes) based on the predicted binding affinity (docking score) and amino acids that are predicted to interact with atoms in the computational fragment. The information output from the FDSL pipeline can then be used to resynthesize the fragments in novel combinations and generate synthetic ligands which could form the basis for potential candidate lead compounds. The FDSL approach thereby contrasts with conventional computational FDBB, which, even when based on predetermined ligand libraries, do not retain information about protein-ligand binding based on initial virtual library screening. For that reason, FDSL constrains the optimization space to identify the best possible trajectories for fragment growth and virtual compound synthesis, and thus can more readily identify promising lead designs.

Here, we expand our FDSL framework introduced in Wilson et al. ^81^ to develop a novel computational drug design methodology that employs two stages of optimization based on applying the fragments as well as the associated fragment attributes derived from ligand prescreening. The two optimization stages proceed as follows: 1) Evolutionary optimization uses principles of natural selection and genetic variation in a computational method for strategically guiding the assembly of synthetic fragments to generate larger compounds. 2) Iterative optimization refines the resulting compounds by adding small fragments to improve bioactivity. At both stages, the fragment information obtained from the FDSL pipeline narrows down the vast chemical space by imposing constraints that limit the search to areas with higher potential for success. Using fragment attributes will both reduce the combinatorial size of the search space and focus the search for ligands on potentially more favorable parts of the space. The resulting computational drug design process not only becomes more efficient, but also more likely to yield compounds with high binding affinities and desirable drug-like properties.

In this paper, we demonstrate the two-staged computational drug design methodology on three distinct protein targets found in different kinds of organisms, i.e., human, bacterial, and viral, and which are in turn implicated in very different kinds of diseases and contexts: 1) Tumor necrosis factor-alpha-induced protein 8-like 2 (TIPE2), a transport protein that can induce leukocyte polarization, sustaining chronic inflammation and ultimately supporting solid cancer tumorigenesis; ^23^ TIPE2 inhibition would provide a therapeutic option for solid tumor cancers. 2) Bacterial protein RelA, which plays a role in detecting amino acid starvation, activating a stringent response in bacteria that leads to persister cell formation. Persister cells can withstand upwards of 1000 times the antibiotic concentrations of their normal cell counterparts; accordingly, inhibiting RelA can allow antibiotics to eradicate bacteria in biofilms. ^31^. 3) the receptor binding domain (RBD) of the S1 subunit of the spike protein (S-protein) of severe acute respiratory syndrome coronavirus 2 (SARS-CoV-2) and the SARS-CoV-2 spike protein receptor binding domain (RBD), which binds to human angiotensin-converting enzyme (ACE-2), thereby facilitating viral entry and representing potential target for antiviral therapeutics for COVID-19. ^78^

The computational studies herein demonstrate the potential of the methods to significantly enhance the efficiency and effectiveness of computational drug design. First, we assess the utility of information about fragments obtained through the initial virtual screening step in FDSL by showing that FDSL with two-stage optimization results can computationally generate candidate ligands that have high predicting binding affinity, with substantially improved performance as compared to not using virtual screening step (i.e., the “naive” approach of conventional computational FBDD). Second, we compare computational ligands generated by FDSL and two-stage optimization to those generated by highly cited and well-documented conventional FBDD methods: 1) Au-toGrow, ^20,73^, which utilizes genetic algorithms, like the first stage optimization of our method; and 2) DeepFrag, ^28,29^ which utilizes deep learning to optimize computational fragment selection and growth based on characteristics of the protein binding pocket, which contrasts to the use of virtual ligand screening based on specific protein targets in our approach. The source code developed to implement the algorithms presented in this paper is available at https://github.com/EESI/FDSL_Evo.

## METHODS

### Ligand Prescreening and Fragmentation Pipeline

The computational drug design procedure begins with prescreening of a ligand library with protein targets. Fig. 1 shows a schematic of the prescreening workflow. The workflow is detailed in previous work, and described in brief here. ^81^ The results presented in this paper used Autodock VINA for prescreening. The crystal structures of the protein files; TIPE2 (PDB ID: TIPE2), RelA (PDB ID: 5IQR), and S-protein (PDB ID: 6M0J); were retrieved from the RCSB Protein Data Bank. The structures were pre-processed by removing waters, co-crystalized proteins, and co-crystalized atoms. Protein structures were prepared in AutoDockTools-1.5.6^68^, including addition of polar hydrogens and calculation of Gasteiger charges. Ligands from an Enamine Ltd. “drug-like” library consisting of around 250,000 molecules were retrieved and optimized using OpenBabel ^56^, generating 3D structures, adding charges and mini-

**Figure 1:**
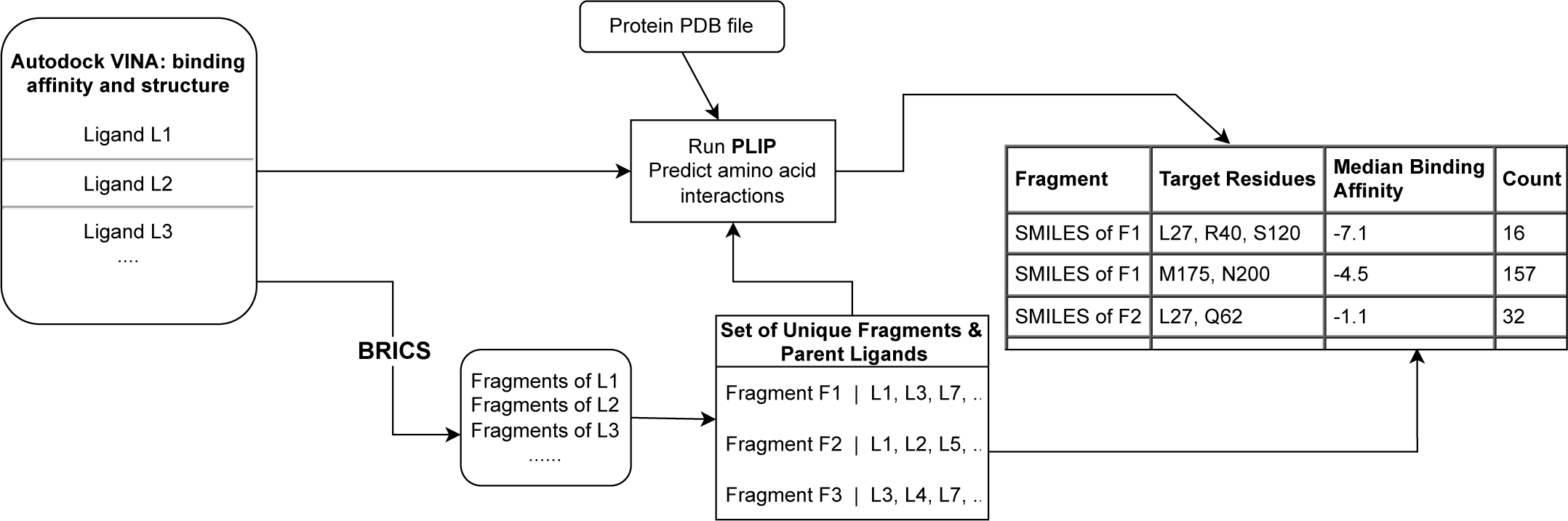
Processing of prescreened ligands begins with the output from prescreening a library of ligands with Autodock VINA, selecting the minimum binding affinity structure. PLIP (Protein-Ligand Interaction Profiler) predicts the binding residues based on the docked ligand-protein complex using a rules-based heuristic algorithm. The ligands are computationally fragmented using the BRICS algorithm. The output from this workflow is a table fragments, identified by their SMILES representation, associated with the binding pocket residues identified by PLIP, as well as the median binding affinity of the parent ligands and the frequency in which the combination is found in the prescreened library. Notably, a fragment will appear multiple times if it is found to bind to different residues (i.e., binding pocket subdomains) in the context of different pre-screened ligands.

### mizing with the MMFF94 force field

High throughput molecular docking calculations of mass libraries were performed using AutoDock Vina 1.2.3 ^21^ on Drexel University Research Computing Facility’s Picotte high performance computing cluster of Intel Xeon Platinum 8268 CPUs. As required for Autodock VINA, grid boxes for ligand docking were generated for each protein targeting known binding sites. The grid box for RelA is centered at x = 297.894, y = 163.593, and z = 219.301 with dimensions of 25.000 Å. For the S-protein the box of dimen-sions 22.000Å *×* 42.000Å *×* 22.000Å were centered at x = −27.878, y = 25.205, and z = 5.514. TIPE2 has a particularly large bind-ing cavity; therefore, 4 grid boxes were generated to span the pocket entrance, thereby occluding the cavity. Consequently, dock-ing with TIPE2 generated 4 times the output, as every ligand was docked in each quadrant grid box. All grid box quadrants are of 12.000Å dimensions with quadrant 1 centered at x = 60.677, y = 5.646, and z = 17.000; quadrant 2 centered at x = 62.636, y = 11.365, and z = 19.959; quadrant 3 centered at x = 68.067, y = 10.738, and z = 18.594; and quadrant 4 centered at x = 67.024, y = 5.362, and z = 17.113.

As shown in Fig. 1, Autodock VINA outputs a PBDQT-formatted file with multiple binding solutions, each with a predicted protein-ligand binding affinity. The lowest binding affinity bound ligand structure is extracted and associated with its binding affinity value. The docked ligand strucure is PDBQT-format file is input to the Protein Ligand Interaction Profiler (PLIP). ^2^ PLIP performs a rule-based prediction of interactions between ligand atoms and protein amino acids, including the bond type and atom–residue pairs. The ligand is also broken into fragments using BRICS, ^15^ an algorithm designed to break bonds in a chemically realistic manner. The fragmentation algorithm is implemented in Python 3.8 using the RDKit open source chemoinformatics software package, available at http://www.rdkit.org. The fragments are then associated with their “parent” ligands (the ligands that were fragmen-tized). Many fragments will have multiple parent ligands, i.e., they will have appeared as the fragments of multiple ligands. The loca-tion of the fragment in the binding pocket is then identified for each parent ligand by finding the maximum common substructure (MCS) between the fragment and ligand. ^76^ Each distinct fragment-subregion combination is then stored, with the fragment stored in a SMILES format ^79^ along with the median binding affinity, the protein residues with bonds identified by PLIP, and the frequency it is found in the prescreened ligand population (i.e., “Count” in Fig. 1).

### Genetic Algorithm (Phase 1 of Fragment Synthesis)

The genetic algorithm requires the target receptor’s PDB (Protein Data Bank) file, an Autodock VINA configuration file (as described in the previous subsection for ligand prescreening), and the output of the fragmentation and analysis pipeline described in the pre-ceding subsection, as shown in Fig. 1. The resulting table is sorted based on the binding affinity to the receptor without considering target residues (i.e. the overall median binding affinity for each fragment). In the current version of the code, BRICS fragments are required, although some modification can make it compatible with any fragmentation protocol.

### Genetic Algorithm Overview

An individual is defined as a collection of fragments used to generate a ligand. Each fragment within this collection is a gene and is represented by its unique index within the source table of library fragments. A rank weighting is calculated and assigned for each index within the source (parent) ligands. These are calculated by the index of the fragments within the input table, and a rank weight-ing function is described below:

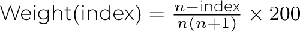

where *n* is the length of the table (number of fragments). The weight index provides larger fractional weights to higher ranked frag-ments towards the top of the list, which are expected to produce ligands of a lower binding affinity because they were generated from ligands with lower binding affinity. These ranked weights are used to define a categorical distribution using the `random.choices()` function, which is used to select indexes when new fragments are being incorporated.

Fig. 2 shows an overview of the genetic algorithm procedure. To start, a random selection of fragments is chosen to seed the first generation. Hydrogens are added to unfilled valences (see Clean Valences block in Fig. 2), and they are run through Autodock VINA to evaluate them (see Autodock Vina block in Fig. 2). In addition, QED scores can be optionally calculated to determine drug like-ness, and a combined QED and VINA score can be used for evaluating candidate ligands in the population instead of VINA score alone in the procedures described below.

**Figure 2:**
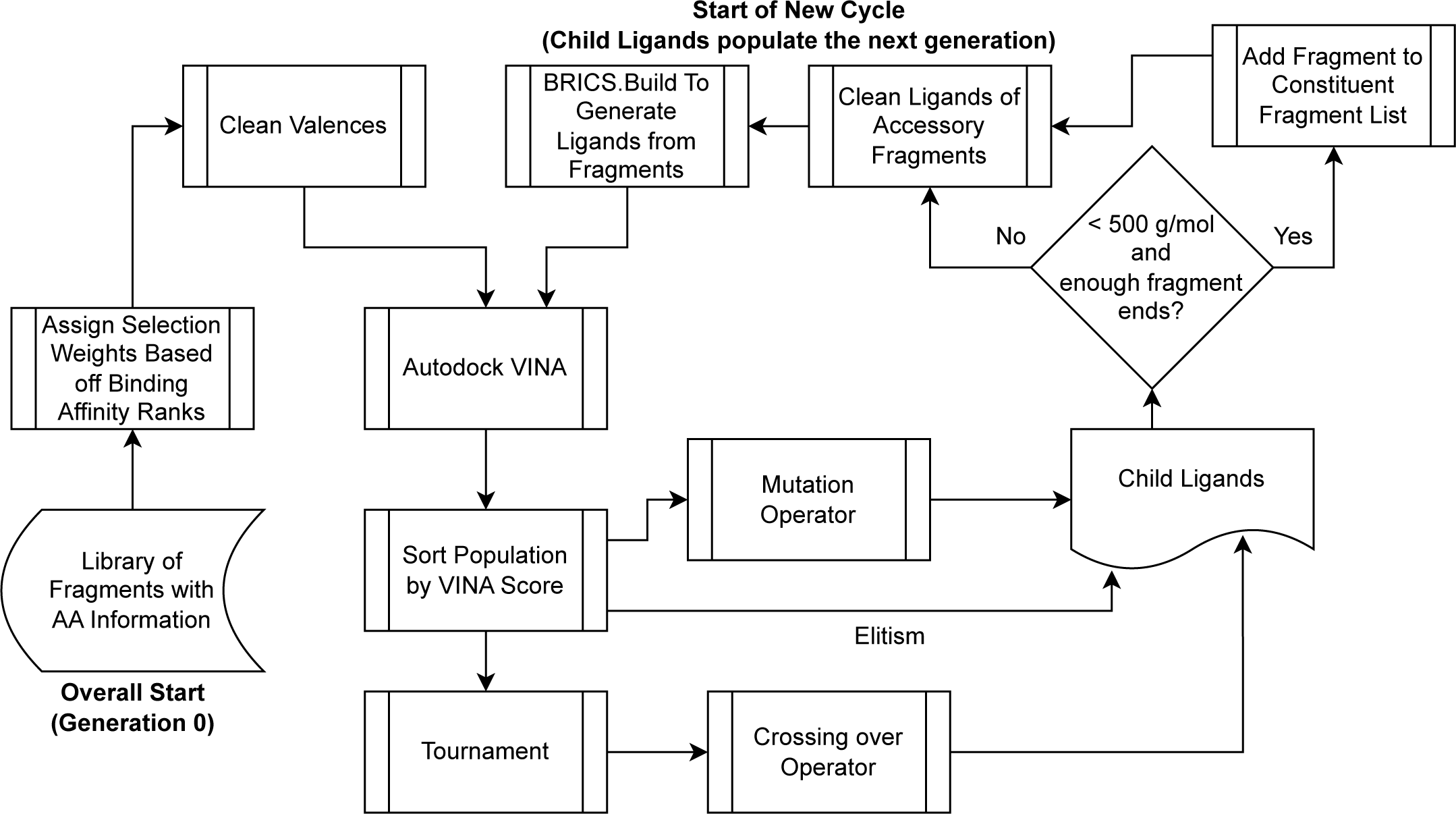
In the first phase of ligand optimization, a genetic algorithm is used to create ligands from the fragments produced by the prescreen-ing and fragmentation pipeline shown in Fig. 1. Fragments are represented like genes and assigned a weighted rank to determine selection probability. Initially, fragments are randomly chosen and evaluated using Autodock VINA (see “Autodock Vina” block) and optionally analyzed for drug likeness via QED scores. Subsequent ligand generations are crafted using mutation (“Mutation” block), crossover, and elitism strategies, abiding by specific molecular weight and fragment use rules. The ligands are further cleaned to ensure all fragments are utilized in the resultant ligand (“Clean Ligands” block) and constructed into new generations using the `BRICS.BUILD` module in RDKit (“`BRICS.BUILD` Generates Ligands” block).

The next generation is determined based off three operators: mutation, crossing over, and elitism. To start, the population is sorted by VINA score (the Sort Population by VINA Score block in Fig. 2), and the top 5/8th of the population are chosen to be ran through the mutation operator (the Mutation block in Fig. 2). These ligands, whether mutated or not, are added to the next generation. Next, a tournament selection takes place (the Tournament block in Fig. 2), where each individual is compared against two other randomly chosen individuals, and the individual with the lowest binding affinity is chosen to be a parent used in the crossing over operator. Two unique individuals who won a tournament are run through the operator at a time, and the operator returns two chil-dren, which will both be included in the next generation. The crossing over operator accounts for 1/4th of the following generation. The last 1/8th of the next generation is developed using elitism, where the individuals with the lowest binding affinity in the parent population are added without alteration.

The children to be used in the next generation are next screened to determine whether they have hit their max molecular weight of 500 g/mol. If not, or the number of fragment ends (unfilled valences) are an odd number, an additional fragment selected based off the rank weighting function described above is added to the ligand (the Add Fragment to Constituent Fragment List block in Fig. 2). If the maximum weight has been reached or the number of fragment ends are even, the ligands remain unaltered and are added to the fragment constituent list and progress to the Clean Ligands phase of the cycle.

To ensure that every fragment included in an individual is incorporated into the resultant ligand, each ligand is ran through a clean-ing function (the Clean Ligands of Accessory Fragments block in Fig. 2) to ensure there are enough fragments ends to accommo-date all fragments. The `BRICS.BUILD` module in RDKit, which is used to generate each ligand from its constituent fragments, will not accommodate a fragment if there is not an end for it to bind to. Therefore, fragments which exceed the number of ends available to attach fragments are pruned, to avoid including it as a gene in future generations when it did not contribute to the evaluated struc-ture. After running through the cleaning function, the `BRICS.BUILD` module generates ligands from the fragments (see `BRICS.BUILD` Generates Ligands from Fragments block in Fig. 2), and the children replace the parents as the new population. Then, the next gen-eration begins.

### Genetic Algorithm Components

#### Mutation Operator

Based off the mutation rate supplied by the user, each fragment has a chance of mutating. In the event of a mutation, another fragment is substituted for the existing fragment, which is selected based off the categorical distribution cal-culated at the beginning using the `random.choices()` function. The ligands, whether mutated or not, are incorporated into the next generation.

#### Crossing Over Operator

The crossing over operator requires two parents to be inputted as well as a user-supplied crossing over rate. If a crossing over occurs, then a random index is selected between both individuals, and the fragments (indexes) between both individuals are swapped after that point. For instance, assume an instance where two individuals with four corresponding frag-ments each are selected as parents:

Parent 1: [314, 132, 4813, 192] Parent 2: [102, 8512, 591, 5123]

The indexes correspond to fragments in the original source CSV. If a crossing over event occurs, a random index is chosen as the in-dex to perform the switch. Suppose the index 2 is selected. The following child ligands will be generated: Child 1: [314, 132, 591, 5123] Child 2: [102, 8512, 4831, 192]

These individuals will be incorporated into the next generation.

#### Generating Ligands from Fragments

The Build function from the BRICS package of RDKIT is used to generate ligands. The func-tion is setup to only output complete SMILES, negating the need to manually add hydrogens to unfilled valences.

### Iterative Fragment Addition (Phase 2 of Fragment Synthesis)

The second stage of optimization is iterative fragment addition, which is effectively a hill-climbing algorithm for maximizing the drug design objective. In this study, we considered both binding affinity alone, as predicted by AutoDock VINA, as well as binding affinity in combination with the Quantitative Effectiveness of Druglikeness (QED) score. The QED score was developed by Bickerton et al. ^8^ as an improvement over rules, such as Lipinski’s Rule of Five ^44^, which combine different thresholds and properties of ligands that tend to be associated with successful drugs. The QED score is a calculated by a formula based on a weighted sum of “desirability functions,” i.e., molecular properties associated with desirability for a particular class of drugs. In this paper, we use the default definition of QED from Bickerton et al., i.e., including molecular weight (MW), octanol-water partition coefficient (ALOGP)24, num-ber of hydrogen bond donors (HBD), number of hydrogen bond acceptors (HBA), molecular polar surface area (PSA), number of rotatable bonds (ROTB), the number of aromatic rings (AROM) and number of structural alerts (ALERTS). ^8^. The QED score is computed using RDKit’s `qed()` module (https://www.rdkit.org/docs/source/rdkit.Chem.QED.html). The algorithm proceeds the same in either the QED + binding affinity or affinity-only, except that for QED + binding affinity, the optimization criterion is a sum of the two scores.

The iterative fragment addition methodology requires a list of premade starter ligands and a dataset including fragments associated with amino acids of the target protein. The list of premade starter ligands is the output of the genetic algorithm, while the dataset of ligands and associated amino acids is the output of the initial prescreening and fragmentation pipeline. Before running the fragment addition, the fragment dataset is converted to a dictionary, where each amino acid is a key with at maximum 100 associated fragments based on the meian binding affinity of parent ligands.

Fig. 3 shows an overview of the iterative fragment addition stage. At the start of each iteration, each ligand in the population is evaluated using Autodock VINA, using the same grid box and seeding as described above for the ligand prescreening stage. Next, the first model in the PDB output file of Autodock VINA is merged with the receptor protein PDB file. This step is in preparation for evaluation of the space using the Protein-Ligand Interaction Profiler (PLIP). Before running PLIP, each ligand in the population is compared against its predecessor to determine if the fragment addition was successful at decreasing binding affinity. If the binding affinity decreased, then the ligand is ready for another attempt at addition. If binding affinity increased, then the addition was unsuccessful at decreasing binding affinity, and the predecessor replaces the current ligand before the addition of a new fragment. Next, all the merged ligand-protein PDB files with molecular weights less than 700 g/mol are ran through PLIP. Ligands with molecular weights greater than 700 g/mol are deemed too large for addition, as larger ligands take increasingly longer to evaluate using VINA and tend to make for worse drug targets. PLIP outputs an XML file containing information about relevant amino acids in the binding pocket of the protein. It also includes distances of the ligand to each of these amino acids. These distances are compared against distances between ligand carbons and protein carbons in the merged ligand-protein PDB, and the closest ligand carbon to an amino acid is selected as the target region to add a fragment.

**Figure 3:**
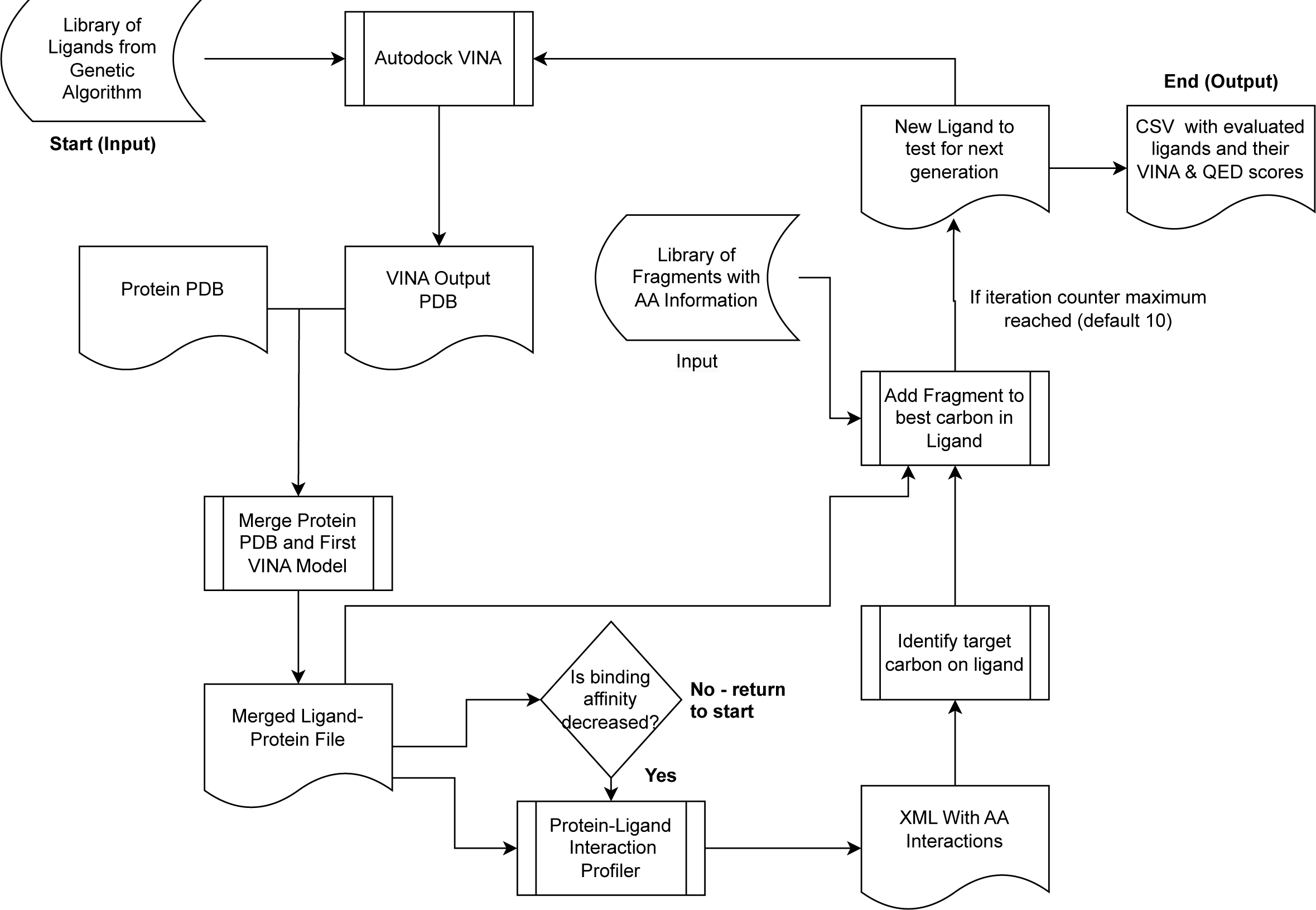
The iterative fragment addition stage, shown schematically here, can begin with any kind of starting ligand but in our method be-gins with candidate ligands synthesized through the previous genetic algorithm-based ligand synthesis phase (see Fig. 2) and fragments ob-tained from the initial ligand prescreening and fragmentation phase (see Fig. 1). The optimization objective of this phase is the binding affinity score predicted by AutoDock VINA, alone or in a sum with the Quantitative Effectiveness of Druglikeness (QED) score, which evaluates benefi-cial molecular properties beneficial for drug design. The methodology begins with premade starter ligands and an amino acid-associated frag-ment dataset. Through successive iterations, each ligand is evaluated and possibly merged with a protein PDB file for further assessment by the Protein-Ligand Interaction Profiler (PLIP). Fragments are strategically added to target regions of the ligand, ensuring optimal binding affinity and maintaining molecular weights under 700 g/mol to ensure viable drug targets. This process cyclically refines ligand structures, using tools like RDKit for optimization and 3D structuring, continuing to a prescribed iteration limit or until an optimizable ligand is generated.

After the target carbon is identified, the ligand and fragment are merged into one MOL object. A bond is formed between the target carbon in the ligand and the atom bound to a dummy atom (indicating a fragment end). All dummy atoms in the fragment are converted to hydrogens to fill the valence of the ligand. The new molecule is embedded into a 3D structure and MMFF94-optimized. In the event RDKit is unable to optimize the newly generated ligand, the next best target carbon is selected, and another fragment addition is attempted. This is repeated multiple times until an optimizable ligand is generated, or the number of iteration attempts reaches ten. If the iteration counter limit is reached, the loop ends, and the SMILES of every ligand and associated VINA and QED score is written to a CSV file. If the iteration counter limit is not reached, the new ligands are evaluated using Autodock VINA and the cycle repeats.

Occasionally, RDKit will be able to embed and optimize the combined ligand and fragment MOL object but will be unable to embed and optimize the SMILES generated from the MOL object. For this reason, a filter is included every generation that tests to make sure that each ligand can be converted from its SMILES to an optimized 3D ligand. SMILES which cannot be converted will not be included in the final CSV file. This prevents inclusion of invalid ligands in the final output which cannot be converted to 3D MOL objects.

## RESULTS

### Assessing Performance of Genetic and Iterative Optimization Ligand Designs Based on Prescreening Information

To determine the effect of fragment quality on the iterative algorithm results, four pools of fragments are created to be fed into the iterative algorithm for each protein target. Each pool is collected from the same prescreening dataset sourced from the fragmenta-tion pipeline illustrated in Fig. 1. The Worst Pool (WP) runs use the worst 1000 fragments by ligand binding affinity. The Large Pool (LP) runs use all fragments, regardless of binding affinity. The Unprioritized (U) and Prioritized (P) runs use the same dataset of frag-ments, where up to 1000 of the best fragments per unique amino acid are included in the pool. The distinction between the two runs is that the P runs pair fragments with interacting amino acids sourced from PLIP. This difference tests the effect of matching fragments with associated amino acids rather than randomly assigning fragments. The WP, LP, and U runs all add random frag-ments within the pool, while the P runs target fragments towards specific amino acids.

The histograms in Fig. 4 highlight the final run results for each protein target. The max ligand size producible by the algorithm is 700 g/mol. Percentile scores and top median pools are highlighted because they tend to be where leads are chosen from. The per-centile scores describe how good the distribution of ligands is towards the top of the results, while the top median scores describe how improved the very best ligands are.

**Figure 4:**
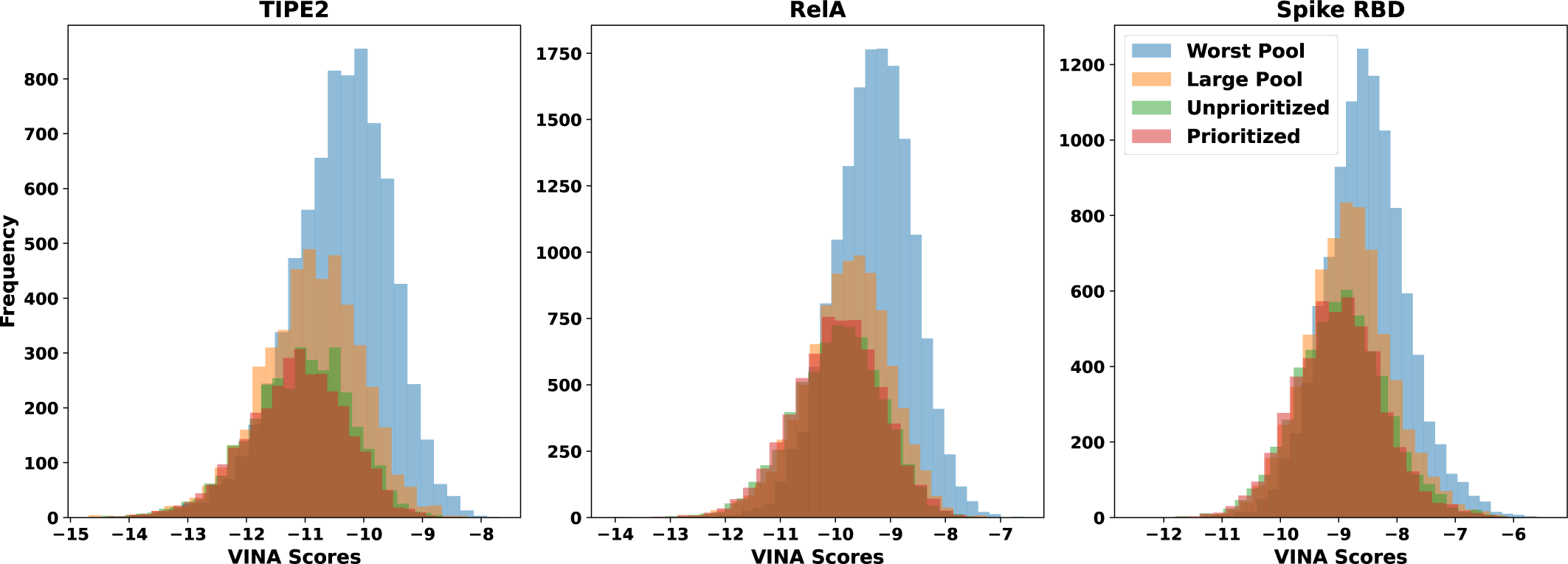
Histogram comparing fragment pool generation methodologies. The top three graphs include comprehensive results of each iterative run. The “Worst Pool” trial used the worst 1000 fragments by VINA score from the source fragment dataset. The “Large Pool” included all fragments. The “Unprioritized” and “Prioritized” trials used the same subset of fragments generated with priority of VINA score and top 1000 fragments associated with a given amino acid. Unprioritized trials use randomly assigned fragments in the pool to bind to the ligands, while Prioritized trials use sub-pools for each amino acid. For a given target amino acid, the Prioritized suggested a fragment known to have interacted with that amino acid in the past based off the PLIP screening.

The P and U plots are shifted left relative to LP and WP runs, indicating a bias towards producing ligands with better binding affinities. For all protein targets, the median and mean VINA scores improve in order of WP, LP, U, and P. In addition, the 95th, 97th, and 99th percentile scores highlight a significant decrease in VINA scores at the top end of each dataset, with significant improvements in VINA scores in the P and U runs relative to the LP and WP runs. However, the P and U runs tend to have percentile scores within ± 0.01 kcal/mol of each other, indicating negligible differences in score between one another. The Top 50 to 10 Median scores for RelA and Spike RBD show a similar pattern, where median scores improve from WP to LP to U/P. Interestingly, the ligands generated for the TIPE2 target did not show a similar trend in score, with LP, U, and P Top 50 to 10 median scores demonstrating no clear trend between runs. The Top 10 Median LP run even outperformed the U and P runs at −14.21 kcal/mol compared to −14.04 kcal/mol and −13.98 kcal/mol respectively. Outlier ligands are often produced by chance. Addition of a fragment to a given carbon may on rare occasions significantly improve binding affinity over targeted efforts to improve the overall distribution of ligands. Therefore, ligands towards the top of the results do not follow the trend of improving binding affinities in the order of WP, LP, and U/P.

Although the Large Pool and Worst Pool runs have lower median binding affinities, they overall produce significantly more ligands relative to the Prioritized and Unprioritized runs, as per the Counts column of each run in Table 1 for each of the protein targets. This is expected as fragments towards the top of the fragment dataset tend to have larger molecular weights. Each addition in the U and P runs increases the molecular higher than an addition in the WP and LP runs, which causes the U and P runs to reach the max molecular weight of 700 g/mol more rapidly. This causes the U and P runs to produce fewer unique ligands than the LP and WP runs.

**Table 1:**
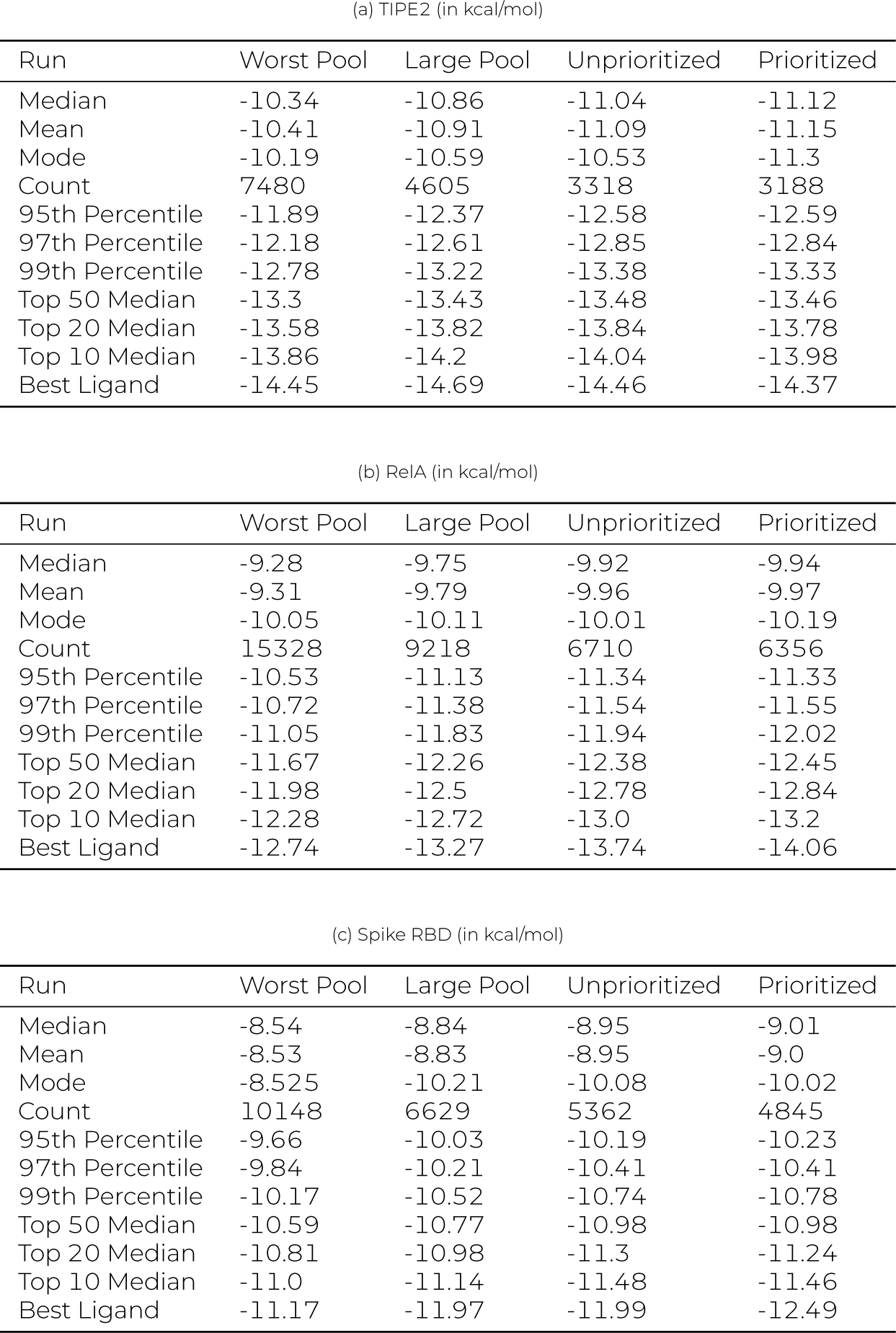
Iterative Run Statistics.

| (a) TIPE2 (in kcal/mol) |  |  |  |  |
| --- | --- | --- | --- | --- |
| Run | Worst Pool | Large Pool | Unprioritized | Prioritized |
| Median | -10.34 | -10.86 | -11.04 | -11.12 |
| Mean | -10.41 | -10.91 | -11.09 | -11.15 |
| Mode | -10.19 | -10.59 | -10.53 | -11.3 |
| Count | 7480 | 4605 | 3318 | 3188 |
| 95th Percentile | -11.89 | -12.37 | -12.58 | -12.59 |
| 97th Percentile | -12.18 | -12.61 | -12.85 | -12.84 |
| 99th Percentile | -12.78 | -13.22 | -13.38 | -13.33 |
| Top 50 Median | -13.3 | -13.43 | -13.48 | -13.46 |
| Top 20 Median | -13.58 | -13.82 | -13.84 | -13.78 |
| Top 10 Median | -13.86 | -14.2 | -14.04 | -13.98 |
| Best Ligand | -14.45 | -14.69 | -14.46 | -14.37 |

| (b) RelA (in kcal/mol) |  |  |  |  |
| --- | --- | --- | --- | --- |
| Run | Worst Pool | Large Pool | Unprioritized | Prioritized |
| Median | -9.28 | -9.75 | -9.92 | -9.94 |
| Mean | -9.31 | -9.79 | -9.96 | -9.97 |
| Mode | -10.05 | -10.11 | -10.01 | -10.19 |
| Count | 15328 | 9218 | 6710 | 6356 |
| 95th Percentile | -10.53 | -11.13 | -11.34 | -11.33 |
| 97th Percentile | -10.72 | -11.38 | -11.54 | -11.55 |
| 99th Percentile | -11.05 | -11.83 | -11.94 | -12.02 |
| Top 50 Median | -11.67 | -12.26 | -12.38 | -12.45 |
| Top 20 Median | -11.98 | -12.5 | -12.78 | -12.84 |
| Top 10 Median | -12.28 | -12.72 | -13.0 | -13.2 |
| Best Ligand | -12.74 | -13.27 | -13.74 | -14.06 |

| (c) Spike RBD (in kcal/mol) |  |  |  |  |
| --- | --- | --- | --- | --- |
| Run | Worst Pool | Large Pool | Unprioritized | Prioritized |
| Median | -8.54 | -8.84 | -8.95 | -9.01 |
| Mean | -8.53 | -8.83 | -8.95 | -9.0 |
| Mode | -8.525 | -10.21 | -10.08 | -10.02 |
| Count | 10148 | 6629 | 5362 | 4845 |
| 95th Percentile | -9.66 | -10.03 | -10.19 | -10.23 |
| 97th Percentile | -9.84 | -10.21 | -10.41 | -10.41 |
| 99th Percentile | -10.17 | -10.52 | -10.74 | -10.78 |
| Top 50 Median | -10.59 | -10.77 | -10.98 | -10.98 |
| Top 20 Median | -10.81 | -10.98 | -11.3 | -11.24 |
| Top 10 Median | -11.0 | -11.14 | -11.48 | -11.46 |
| Best Ligand | -11.17 | -11.97 | -11.99 | -12.49 |

### Comparison of Prescreening and Optimization Methodology with Other Genetic and Deep Learning Methods

In addition to the above experiments, trials of other ligand optimization packages are shown here to compare the effectiveness of the described state-of-art genetic and iterative (machine learning) methodologies. One exemplary package that takes a genetic approach is Autogrow4. Due to limitations in ligand pool size, for the trials shown here, Autogrow4 is fed a random sample of 1000 source ligands from the top 10000 ligands used by the fragmentation pipeline and was ran 10 times. All default variables and pack-ages were used, and the file conversion package selected is `obabel`. Each run is allowed to run for 30 generations, at which scores between generations tend to plateau. The same receptors and search boxes used by the genetic and iterative code are supplied to Autogrow4.

The histograms in Fig. 5 highlight significantly higher binding affinities for ligands generated by Autogrow4 relative to binding affinities generated by the iterative methodology. This is further highlighted in the tables, which show significant improvements in binding affinity in the percentile score columns and the top ligand median score columns. Interestingly, Autogrow4 appeared to generate a better overall median score in the RelA trial. The best ligands produced by Autogrow4 in all 10 runs for each target pro-tein had VINA scores of −12.6, −11.8, and −10.3 kcal/mol for TIPE2, RelA, and Spike RBD respectively, while the best iterative ligands from the Prioritized runs were −14.37, −14.06, and −12.49 kcal/mol.

**Figure 5:**
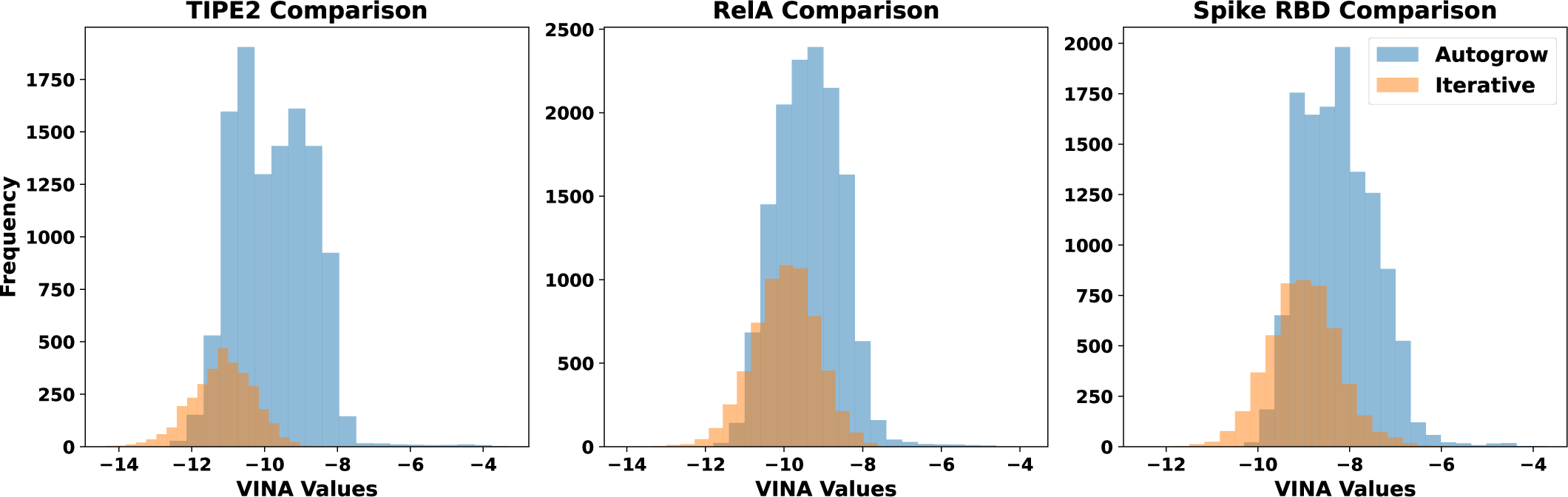
Comparison of Autogrow4 and Iterative generated ligands. The proposed method’s histogram data was selected from Prioritized run described in Fig. 4 and accompanying text.

The second-stage iterative optimization of the proposed approach appears to produce significantly fewer ligands than Autogrow4, as indicated by the counts in Table 2. This is attributable to the iterative approach only being able to add fragments onto ligands, which means it will hit the molecular weight ceiling faster than an exclusively genetic approach like Autogrow4. Autogrow4 can mix and match substructures within a given population of molecules, allowing for more combinations and hence more unique lig-ands.

**Table 2:** AutoGrow4 Comparison (in kcal/mol)

| Run | TIPE2 |  | RelA |  | Spike RBD |  |
| --- | --- | --- | --- | --- | --- | --- |
|  | Autogrow4 | Prioritized | Autogrow4 | Prioritized | Autogrow4 | Prioritized |
| Median | -9.8 | -11.12 | -9.4 | -9.94 | -8.3 | -9.01 |
| Mean | -9.77 | -11.15 | -9.41 | -9.97 | -8.25 | -9.0 |
| Mode | -10.7 | -11.3 | -9.1 | -10.19 | -8.5 | -10.02 |
| Count | 11142 | 3188 | 13760 | 6356 | 12215 | 4845 |
| 95th Percentile | -11.3 | -12.59 | -10.7 | -11.33 | -9.4 | -10.23 |
| 97th Percentile | -11.5 | -12.84 | -10.8 | -11.55 | -9.5 | -10.41 |
| 99th Percentile | -11.8 | -13.33 | -11.1 | -12.02 | -9.7 | -10.78 |
| Top 50 Median | -12.2 | -13.46 | -11.4 | -12.45 | -9.9 | -10.98 |
| Top 20 Median | -12.4 | -13.78 | -11.6 | -12.84 | -10.0 | -11.24 |
| Top 10 Median | -12.45 | -13.98 | -11.7 | -13.2 | -10.1 | -11.46 |
| Best Ligand | -12.6 | -14.37 | -11.8 | -14.06 | -10.3 | -12.49 |

The proposed approach is also compared to DeepFrag, a deep learning approach which aims to predict the best fragment to add to a ligand within a binding pocket. The default fragments in the DeepFrag library are used in this comparison, and 10 runs are com-pleted for each target. To create a fair comparison, each iteration, the best ligand proposed by DeepFrag is used as the input ligand for the next iteration. 10 iterations are completed per run. The intermediate ligands produced by DeepFrag are combined into one dataset which is sorted by DeepFrag scoring function. Due to computational constraints, a sample of the best ten thousand ligands from this combined dataset are ran through Autodock VINA. A histogram comparison would be ineffective at comparing the en-tire population of ligands generated by the proposed methodology to a sample of the DeepFrag results, so a bar plot is used to to represent the results in Fig. 6.

**Figure 6:**
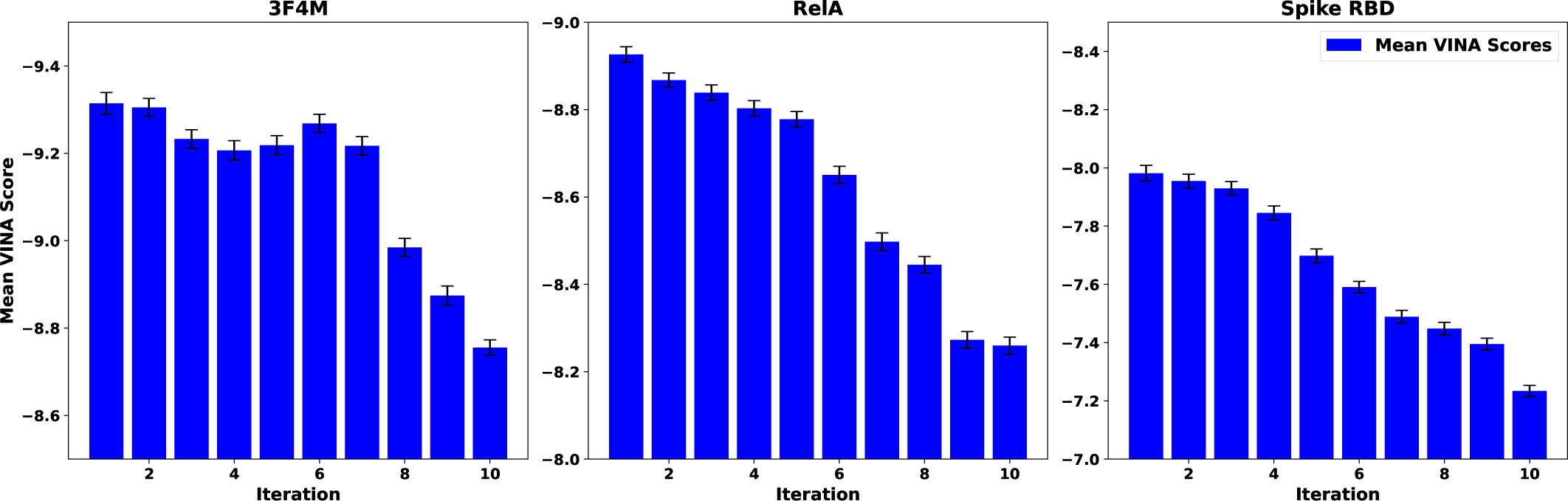
The mean VINA score of each iteration in the Deep Frag runs is plotted. The unbiased standard error of the mean is calculated for each iteration and displayed as error bars. Scores tend to increase per iteration of DeepFrag, indicating worsened binding affinities. The best VINA scores for each protein target are −12.17, −11.47, and −9.895 kcal/mol for TIPE2, RelA, and Spike RBD respectively. The graph is plotted such that the y-axis values decrease from bottom to top to show stronger binding affinities as higher values. Also, the y-axis range differs for each target to illustrate the similarity in trend between the targets differently for each target.

The trend in the bar plots in Fig. 6 show that DeepFrag appears to produce poor ligands for the target binding pockets. Even when comparing best scores, DeepFrag produces significantly worse scores than the best scores of the proposed iterative methodol-ogy, at −14.37, −14.06, and −12.49 kcal/mol for TIPE2, RelA, and Spike RBD respectively. Interestingly, the average scores tend to trend down, indicating that DeepFrag may not be optimized to improve binding affinity generated by Autodock VINA. This may be caused by DeepFrag being over-fitted to the training set used, limiting extension to other protein targets like the ones presented in this study. Because the model is not tuned for the tested protein targets, the ligands generated drift into higher Autodock VINA scores, indicating decreased (poorer) binding affinities.

### Evaluating Multi-Objective Optimization for Druglikeness and Binding Affinity Based on Prescreening Information

To show that the proposed method can also account for drug likeness properties in addition to binding affinity, and still produce viable candidate ligands, a straightforward modification can be made to render the optimization multiobjective. Specifically, in ad-dition to evaluating binding affinity using Autodock VINA, the genetic and iterative algorithms contain an optional multi-objective scoring function. The multi-objective scoring function considers QED score, a single metric generated from drug desirability func-tions, in addition to VINA binding scores. This scoring function can be customized depending on a user’s optimization interests. The algorithm combines a ligand’s VINA and QED z-scores into a single value, giving equal weight to both. The z-scores are calculated relative to the original ligand dataset from which the fragments were sourced. This methodology ensures that marginal gains in VINA performance do not significantly reduce drug likeness.

The graphs in Fig. 7 compare the generated ligands using the multi-objective approach compared to a VINA prioritization approach. Both approaches are ran on the same set of starter ligands generated from the genetic algorithm, which is ran using the multi-objective evaluation function. Table 3 summarizes the statistics for each run for the respective protein targets.

**Figure 7:**
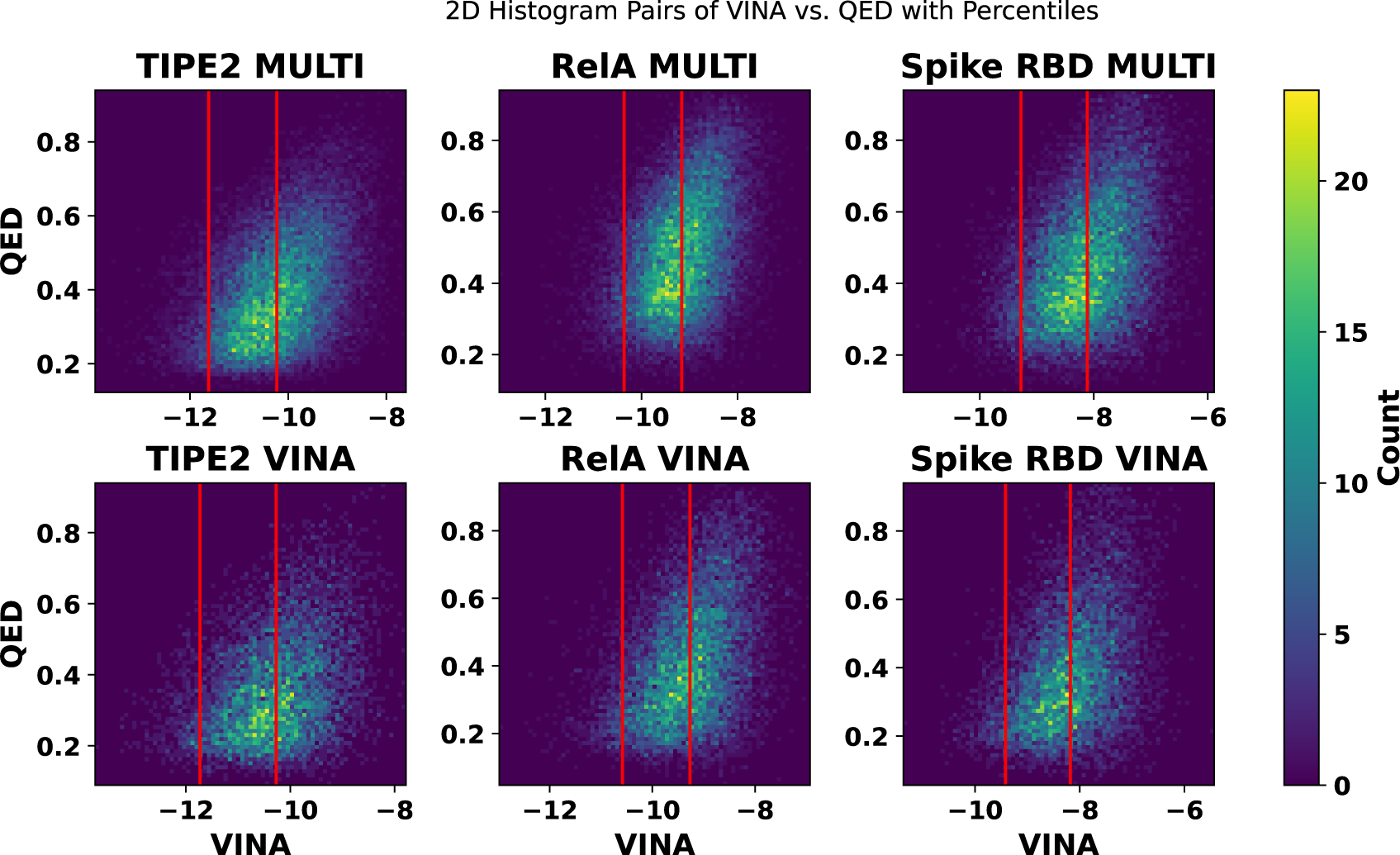
Histograms comparing multi-objective and VINA prioritizations over final VINA and QED scores using iterative approach. Lower VINA scores indicate improved binding affinity and higher QED scores indicate better drug-likeness. Red lines indicate 50th and 95th percentile scores, which are selected to segment regions of each dataset. Both iterative runs are run on the same set of starting ligands from the genetic algorithm, which is run with multi-objective prioritization. The starting ligands are selected based on best multi-objective score.

**Table 3:** Multiobjective Iterative Run Statistics (in kcal/mol)

| Run | TIPE2 |  | RelA |  | Spike RBD |  |
| --- | --- | --- | --- | --- | --- | --- |
|  | MULTI | VINA | MULTI | VINA | MULTI | VINA |
| Median | -10.23 | -10.27 | -9.16 | -9.27 | -8.12 | -8.18 |
| Mean | -10.24 | -10.29 | -9.17 | -9.3 | -8.12 | -8.19 |
| Mode | -10.22 | -10.19 | -10.01 | -10.07 | -8.177 | -8.398 |
| Count | 24748 | 7784 | 24083 | 11886 | 19440 | 9041 |
| 95th Percentile | -11.63 | -11.73 | -10.36 | -10.58 | -9.28 | -9.42 |
| 97th Percentile | -11.83 | -11.94 | -10.55 | -10.77 | -9.43 | -9.62 |
| 99th Percentile | -12.25 | -12.31 | -10.9 | -11.17 | -9.71 | -9.91 |
| Top 50 Median | -13.0 | -12.73 | -11.57 | -11.73 | -10.2 | -10.28 |
| Top 20 Median | -13.27 | -12.99 | -11.87 | -12.03 | -10.39 | -10.42 |
| Top 10 Median | -13.5 | -13.16 | -12.0 | -12.29 | -10.62 | -10.56 |
| Best Ligand | -13.93 | -13.74 | -12.96 | -12.98 | -11.34 | -11.37 |

The plots in Fig. 7 demonstrate that the multi-objective runs produces similar VINA score distributions to the VINA prioritization runs. Although the multi-objective prioritization produces slightly worse percentile scores, it produced better Top 50, 20, and 10 median scores during the TIPE2 runs and a better Top 10 median score during the Spike RBD run, as shown in Table 3). Fig. 8 indi-cates that the multi-objective function produces ligands with similar binding affinities to the VINA prioritization, while significantly improving QED scores, even at the top of the datasets where VINA scores tend to be improved but QED scores tend to be lower.

**Figure 8:**
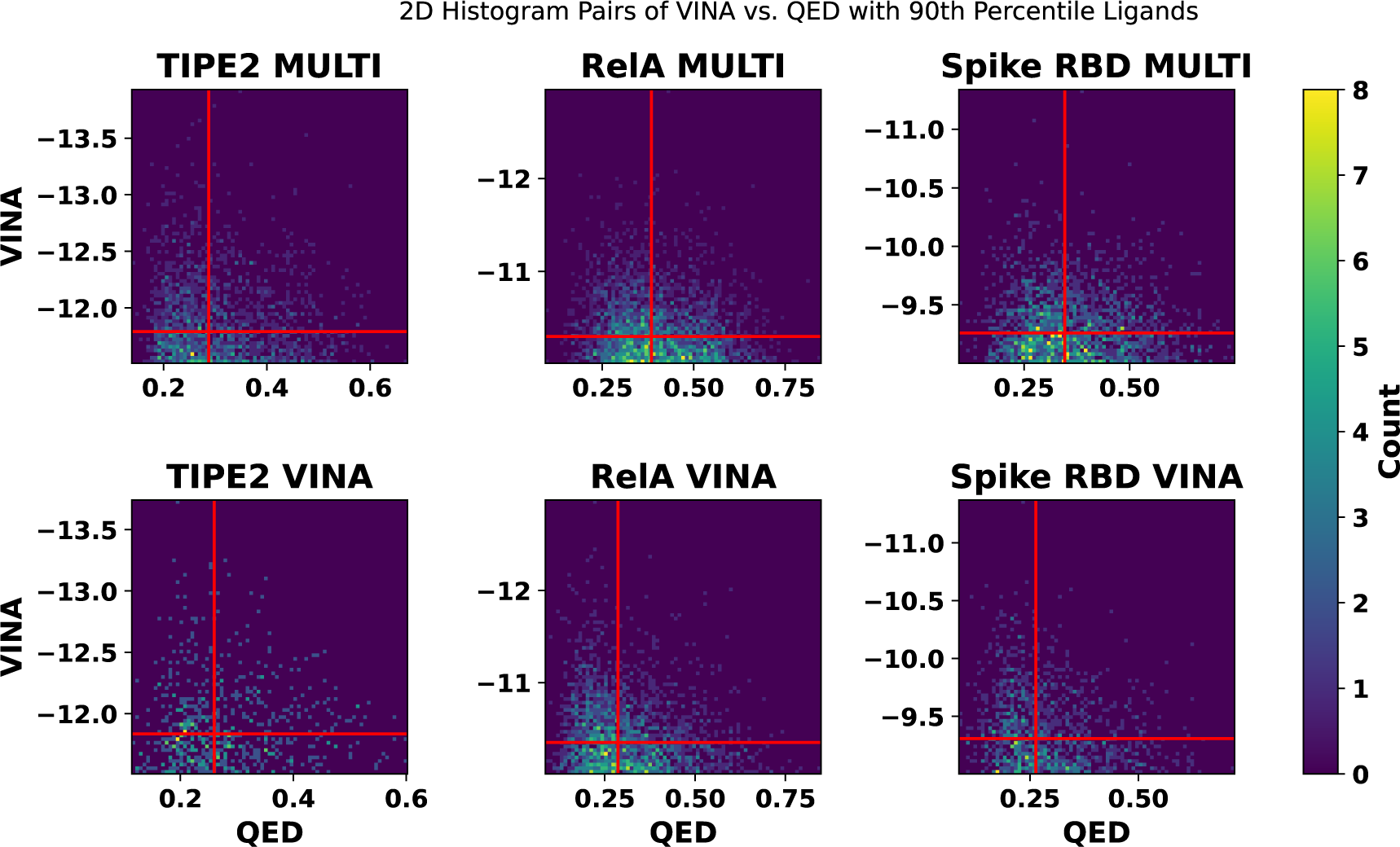
Focusing on the strongest ligand candidates, with respect to predicted binding affinity, the histograms shown here plot the 95th percentile ligands by VINA score with QED score for each of the protein targets. Horizontal line indicates 97.5th percentile VINA score. Vertical line indicates the 50th percentile QED score of the 95th percentile ligands by VINA score.

Similar to the Large Pool and Worst Pool trials described above, the multi-objective runs produced far more ligands than the VINA prioritization runs, for example, as indicated by the counts in Table 3. However, both trials used the same fragment pools. The rea-son for this difference is that the multi-objective trials account for drug desirability, which tends to prefer smaller ligands. If a given iteration produces a marginal gain in VINA score but a large gain in molecular weight, the algorithm will backtrack as the multi-objective score will not have improved, even if the VINA score did. Therefore, more unique ligands are produced as each iteration needs to improve VINA score significantly enough to outweigh any decreases in QED score.

### Identifying Structures for Potential Candidate Ligands

To illustrate the kinds of structures that are produced as a result of the proposed pipeline and optimization methods, the struc-tural formula and SMILES strings of three potential candidate ligands for each of the targets analyzed in this paper (TIPE2, RelA, and Spike RBD) are shown in Table 4. These ligands, for example, may be evaluated in future *in vitro* studies for binding affinity and druggability. The candidate structures shown here are selected from the results of both the default-objective and multi-objective 2D Histogram Pairs of VINA vs. QED with Percentiles functions described in previous sections. The criteria used to select potential exemplary candidates to display in Table 4 are (i) to minimize binding affinity as predicted by Autodock VINA (specifically targeting predicted affinities of less than −13 kcal/mol or as close as possible where targets were not found in that range), (ii) estimated solubility (ESOL) scores indicating moderate solubility or better calculated to be between −4 and −6 using the method described in ^16^, and (iii) molecular weights of less than 700 g/mol. Additional quantitative values for drug-likeness properties are predicted by SwissADME ^13^ and shown in Table 4.

**Table 4:** Molecular Structures of Potential Candidate Ligands Generated by the FDSL and Two-Stage Optimization Pipeline.

|  | Binding Affinity (kcal/mol) | MW (g/mol) | ESOL | MLogP | TPSA (Å²) | Log Kp (cm/s) | GI | BBB |
| --- | --- | --- | --- | --- | --- | --- | --- | --- |
| (i) | -13.04 | 557.64 | -5.82 | 5.28 | 70.56 | -6.76 | High | Yes |
| (ii) | -12.85 | 555.65 | -5.81 | 4.07 | 97.24 | -7.14 | High | No |
| (iii) | -12.40 | 602.72 | -5.77 | 3.42 | 78.09 | -7.44 | High | No |
| (iv) | -12.83 | 625.68 | -5.60 | 4.59 | 129.87 | -7.97 | High | No |
| (v) | -12.29 | 584.67 | -4.30 | 3.41 | 124.32 | -8.87 | High | No |
| (vi) | -12.18 | 548.55 | -5.41 | 2.51 | 129.03 | -7.42 | High | No |
| (vii) | -11.40 | 597.62 | -4.61 | 2.69 | 178.92 | -8.75 | Low | No |
| (viii) | -11.24 | 676.60 | -4.47 | 0.43 | 237.15 | -9.93 | Low | No |
| (ix) | -11.12 | 552.54 | -5.65 | 4.94 | 135.21 | -7.55 | Low | No |

## DISCUSSION

As the results presented in this paper illustrate, the Fragments from Ligands Drug Discovery (FDSL) pipeline can significantly improve computational ligand design and optimization by prioritizing fragments based on the results of initial virtual screening. By prioritizing fragments with higher potentials for success, the FDSL pipeline not only increases the efficiency of the subsequent drug design process but also it towards yielding compounds with optimal binding affinities and drug-like properties. Moreover, the FDSL pipeline includes ligand-binding domain analysis, such that key carbons for binding are identified and prioritized during the iterative step, thereby improving optimization as well.

Specifically, the results show that fine-tuned fragment pools, including fragments involving ligands with lower binding affinities, produce larger shares of ligands with good binding affinities relative to pools without prioritization. Additionally, the similar perfor-mance of the Prioritized trials (fragment pools associated by amino acid) and Unprioritized trials (Prioritized fragment pools without amino acid association) hints at a minimal influence of specifying fragments to specific amino acids during the fragment addition process. Outlier ligands with significant performance often emerge due to chance, which blurs the observable trends at the top tier of each resulting dataset. This can be attributed to fragments within the Large Pool (pool with all fragments) and Worst Pool (pool with worst 1000 fragments by binding affinity) that significantly improve binding affinity in contexts not apparent in the source lig- and dataset. Furthermore, fragments included in the Prioritized and Unprioritized pools with higher associated binding affinities tend to be heavier, hitting the maximum molecular weight of 700 g/mol in fewer iterations and thus producing fewer unique lig-ands than the Large Pool and Worst Pool runs. This upper limit is selected to avoid limiting max ligand sizes to 500 g/mol accord-ing to Lipinski’s Rule of Five, which, as others have pointed out, are likely outdated and, if used strictly, will filter out potential solid leads ^70^. Raising the upper limit beyond 700 g/mol may result in ligands displaying stronger binding affinities. These ligands would be better suited for the large binding pockets of the target proteins under consideration. However, it is important to note that this limit is chosen to enhance computational efficiency. Using larger ligands in Autodock VINA significantly increases the computa-tional time required, which may limit the number of unique ligands screened within a given timeframe.

The proposed method is tested on three distinct types of targets, each with its unique challenges. First, we tested protein targets associated with solid cancer tumorigenesis, aiming for better cancer treatments. Our second design target relates to the issue of antimicrobial resistance, a significant concern as rising resistance could render basic infections untreatable. Third, we applied the proposed method to the spike protein Receptor Binding Domain (RBD) of COVID−19, where inhibiting this domain might prevent the virus from entering human cells, offering a potential therapeutic avenue. The robustness of this approach is further reinforced by its capacity to handle the varied geometries, binding conditions, and chemical conditions presented by these targets. Navigat-ing different geometries means understanding diverse target structures, while adapting to unique binding conditions and varying chemical environments showcases the method’s versatility.

The methodology yields ligands with enhanced binding affinities when compared to other methods. This improvement may be due to the capability of the genetic and iterative algorithms adapted for the FDSL pipeline in this paper to accommodate larger source ligand datasets. In contrast, Autogrow4 and DeepFrag have limitations in terms of source ligand dataset size. The scalability of the proposed methodology enables more extensive exploration and analysis of the chemical space within the binding pocket prior to ligand generation. Scalability, in turn, makes it possible for the genetic and iterative algorithms with an even greater amount of information for constructing new ligands and fine-tuning them towards improved binding affinities than demonstrated in this paper.

Balancing optimal binding affinity with favorable drug-likeness properties is essential for successful lead identification in drug discovery. The method as currently developed employs Quantitative Estimate of Druglikeness (QED) scores. The results demonstrate that employing a multi-objective evaluation can produce candidate leads that can have superior druglikeness properties, such as solubility, with strong binding affinity. In particular, multi-objective prioritization produces a more diverse pool of ligands compared to VINA prioritization. The generated ligands demonstrate significant improvements in QED scores with minimal losses in binding affinity. Moreover, by prioritizing QED scores, modest improvements in binding affinity do not override significant decreases in drug-likeness, resulting in ligands with *both* improved binding affinities and drug-likeness.

Notably, the leads in Table 4 have ESOL scores between −4 and −6, indicating moderate solubility. ^16^ To prepare for *in vitro* studies, polar groups may be manually added to further improve solubility, though these changes may affect predicted binding affinity. Other computational estimations of solubility, such as MLogP, can also be assessed to determine if log P scores are less than 5, which is in agreement with Lipinski’s rule of fives. ^44^

Although the Prioritized runs did not yield significant improvements in binding affinity relative to the Unprioritized runs, further pool screening should be completed in future studies to determine if amino acid matching can yield stronger ligands. For instance, matching fragment properties to amino acid properties instead of exclusively relying on PLIP analysis may yield more optimal fragment-amino acid pairings, improving ligand binding affinity upon addition. Future studies may also consider adding other objectives for optimization.

Finally, the proposed method relies on *computationally predicted* rather than actual experimental data on the initial ligand population— while still providing useful information to guide the drug design and optimization process. Not only is the prescreening data generated *in silico*, thereby avoiding costs and complexity of *in vitro* screening, but the prescreening is also done using relatively low computational-cost and scalable computer docking methods, as opposed to more costly and less scalable molecular dynamics methods. Accordingly, the FDSL and optimization methods presented in this paper are highly scalable. To further scalability, the code developed to implement both the genetic and iterative optimization stages herein is fully parallelized, and can be readily executed across multiple processors in a computing environment as the ligand population and fragment pool increases.

## DATA AND SOFTWARE AVAILABILITY

The source code for the optimization methods described in this paper has been made publicly available for non-commercial use only at https://github.com/EESI/FDSL_Evo. The scripts used for running the trials shown in this paper are also available at the Github, and they can provide a guide for implementation in other high performance computing environments. The files used as inputs to the optimization pipeline and raw results of the optimized pipeline shown in this paper are also provided at the same Github site.

## ACKNOWLEDGMENTS

The work presented here is the result of code developed and run using high performance computing resources through Drexel’s University Research Computing Facility. This work was partially supported by the NSF REU supplement to grant #1919691, as well as NSF grants #1936791, #1936782 and #2107108.

## REFERENCES

[1] S. Abouchekeir, A. Vu, M. Mukaidaisi, K. Grantham, A. Tchagang, and Y. Li. Adversarial deep evolutionary learning for drug design. Bio Systems, 222:104790, Dec. 2022.

[2] M. F. Adasme, K. L. Linnemann, S. N. Bolz, F. Kaiser, S. Salentin, V. J. Haupt, and M. Schroeder. PLIP 2021: Expanding the scope of the protein–ligand interaction profiler to DNA and RNA. Nucleic Acids Research, 49(W1):W530–W534, July 2021.

[3] S. Ahn, J. Kim, H. Lee, and J. Shin. Guiding Deep Molecular Optimization with Genetic Exploration. In *Advances in Neural Information Processing Systems*, volume 33, pages 12008–12021. Curran Associates, Inc., 2020.

[4] W. J. Allen, B. C. Fochtman, T. E. Balius, and R. C. Rizzo. Customizable de novo Design Strategies for DOCK: Application to HIVgp41 and Other Therapeutic Targets. Journal of computational chemistry, 38(30):2641–2663, Nov. 2017.

[5] J. Arús-Pous, A. Patronov, E. J. Bjerrum, C. Tyrchan, J.-L. Reymond, H. Chen, and O. Engkvist. SMILES-based deep generative scaffold decorator for de-novo drug design. Journal of Cheminformatics, 12(1):38, May 2020.

[6] Y. Bian and X.-Q. Xie. Generative chemistry: Drug discovery with deep learning generative models. Journal of Molecular Modeling, 27(3):71, Feb. 2021.

[7] Y. Bian and X.-Q. S. Xie. Computational Fragment-Based Drug Design: Current Trends, Strategies, and Applications. The AAPS journal, 20(3):59, Apr. 2018.

[8] G. R. Bickerton, G. V. Paolini, J. Besnard, S. Muresan, and A. L. Hopkins. Quantifying the chemical beauty of drugs. Nature Chemistry, 4(2):90–98, Jan. 2012.

[9] T. Blaschke, M. Olivecrona, O. Engkvist, J. Bajorath, and H. Chen. Application of generative autoencoder in de novo molecular design, Nov. 2017.

[10] M. Bon, A. Bilsland, J. Bower, and K. McAulay. Fragment-based drug discovery-the importance of high-quality molecule libraries. Molecular Oncology, 16(21):3761–3777, Nov. 2022.

[11] N. Brown, M. Fiscato, M. H. Segler, and A. C. Vaucher. GuacaMol: Benchmarking Models for de Novo Molecular Design. Journal of Chemical Information and Modeling, 59(3):1096–1108, Mar. 2019.

[12] Á. Cortés-Cabrera, F. Gago, and A. Morreale. A computational fragment-based de novo design protocol guided by ligand efficiency indices (LEI). Methods in Molecular Biology (Clifton, N.J.), 1289:89–100, 2015.

[13] A. Daina, O. Michielin, and V. Zoete. SwissADME: A free web tool to evaluate pharmacokinetics, drug-likeness and medicinal chemistry friendliness of small molecules. Scientific Reports, 7(1):42717, Mar. 2017.

[14] L. R. de Souza Neto, J. T. Moreira-Filho, B. J. Neves, R. L. B. R. Maidana, A. C. R. Guimarães, N. Furnham, C. H. Andrade, and F. P. Silva. In silico Strategies to Support Fragment-to-Lead Optimization in Drug Discovery. Frontiers in Chemistry, 8, 2020.

[15] J. Degen, C. Wegscheid-Gerlach, A. Zaliani, and M. Rarey. On the art of compiling and using ’drug-like’ chemical fragment spaces. ChemMedChem, 3(10):1503–1507, Oct. 2008.

[16] J. S. Delaney. ESOL: Estimating aqueous solubility directly from molecular structure. Journal of Chemical Information and Computer Sciences, 44(3):1000–1005, 2004.

[17] F. Dey and A. Caflisch. Fragment-Based de Novo Ligand Design by Multiobjective Evolutionary Optimization. Journal of Chemical Information and Modeling, 48(3):679–690, Mar. 2008.

[18] B. C. Doak and J. Kihlberg. Drug discovery beyond the rule of 5 -Opportunities and challenges. Expert Opinion on Drug Discovery, 12(2):115–119, Feb. 2017.

[19] H. Dowden and J. Munro. Trends in clinical success rates and therapeutic focus. Nature Reviews Drug Discovery, 18(7):495– 496, May 2019.

[20] J. D. Durrant, R. E. Amaro, and J. A. McCammon. AutoGrow: A Novel Algorithm for Protein Inhibitor Design. Chemical Biology & Drug Design, 73(2):168–178, 2009.

[21] J. Eberhardt, D. Santos-Martins, A. F. Tillack, and S. Forli. AutoDock Vina 1.2.0: New Docking Methods, Expanded Force Field, and Python Bindings. Journal of Chemical Information and Modeling, 61(8):3891–3898, Aug. 2021.

[22] D. A. Erlanson, S. W. Fesik, R. E. Hubbard, W. Jahnke, and H. Jhoti Twenty years on: The impact of fragments on drug discovery. Nature Reviews Drug Discovery, 15(9):605–619, Sept. 2016.

[23] S. A. Fayngerts, Z. Wang, A. Zamani, H. Sun, A. E. Boggs, T. P. Porturas, W. Xie, M. Lin, T. Cathopoulis, J. R. Goldsmith, A. Vourekas, and Y. H. Chen. Direction of leukocyte polarization and migration by the phosphoinositide-transfer protein TIPE2. Nature Immunology, 18(12):1353–1360, Dec. 2017.

[24] N. C. Firth, B. Atrash, N. Brown, and J. Blagg. MOARF, an Integrated Workflow for Multiobjective Optimization: Implementation, Synthesis, and Biological Evaluation. Journal of Chemical Information and Modeling, 55(6):1169–1180, June 2015.

[25] J. C. Fromer and C. W. Coley. Computer-aided multi-objective optimization in small molecule discovery. Patterns, 4(2), Feb. 2023.

[26] K. Grantham, M. Mukaidaisi, H. K. Ooi, M. S. Ghaemi, A. Tchagang, and Y. Li. Deep Evolutionary Learning for Molecular Design. IEEE Computational Intelligence Magazine, 17(2):14–28, May 2022.

[27] D. Grechishnikova. Transformer neural network for protein-specific de novo drug generation as a machine translation problem. Scientific Reports, 11(1):321, Jan. 2021.

[28] H. Green and J. D. Durrant. DeepFrag: An Open-Source Browser App for Deep-Learning Lead Optimization. Journal of Chemical Information and Modeling, 61(6):2523–2529, June 2021.

[29] H. Green, D. R. Koes, and J. D. Durrant. DeepFrag: A deep convolutional neural network for fragment-based lead optimization. Chemical Science, 12(23):8036–8047, May 2021.

[30] R. Gupta, D. Srivastava, M. Sahu, S. Tiwari, R. K. Ambasta, and P. Kumar. Artificial intelligence to deep learning: Machine intelligence approach for drug discovery. Molecular Diversity, 25(3):1315–1360, Aug. 2021.

[31] L. Hall-Stoodley, L. Nistico, K. Sambanthamoorthy, B. Dice, D. Nguyen, W. J. Mershon, C. Johnson, F. Z. Hu, P. Stoodley, G. D. Ehrlich, and J. C. Post. Characterization of biofilm matrix, degradation by DNase treatment and evidence of capsule downregulation in Streptococcus pneumoniae clinical isolates. BMC microbiology, 8:173, Oct. 2008.

[32] R. K. Harrison. Phase II and phase III failures: 2013–2015. Nature Reviews Drug Discovery, 15(12):817–818, Dec. 2016.

[33] S. Honda, S. Shi, and H. R. Ueda. SMILES Transformer: Pre-trained Molecular Fingerprint for Low Data Drug Discovery, Nov. 2019.

[34] C.-L. Hung and C.-C. Chen. Computational Approaches for Drug Discovery. Drug Development Research, 75(6):412–418, 2014.

[35] S. A. Jacobs, T. Moon, K. McLoughlin, D. Jones, D. Hysom, D. H. Ahn, J. Gyllenhaal, P. Watson, F. C. Lightstone, J. E. Allen, I. Karlin, and B. Van Essen. Enabling rapid COVID−19 small molecule drug design through scalable deep learning of generative models. The International Journal of High Performance Computing Applications, 35(5):469–482, Sept. 2021.

[36] A. Kerstjens and H. De Winter. LEADD: Lamarckian evolutionary algorithm for de novo drug design. Journal of Cheminformatics, 14(1):3, Jan. 2022.

[37] P. Kirsch, A. M. Hartman, A. K. H. Hirsch, and M. Empting. Concepts and Core Principles of Fragment-Based Drug Design. Molecules, 24(23):4309, Nov. 2019.

[38] A. Kruel, A. McNaughton, and N. Kumar. Scaffold-Based Multi-Objective Drug Candidate Optimization, Dec. 2022.

[39] Y. Kwon and J. Lee. MolFinder: An evolutionary algorithm for the global optimization of molecular properties and the extensive exploration of chemical space using SMILES. Journal of Cheminformatics, 13(1):24, Mar. 2021.

[40] Q. Li. Application of Fragment-Based Drug Discovery to Versatile Targets. Frontiers in Molecular Biosciences, 7:180, 2020.

[41] Y. Li, J. Pei, and L. Lai. Structure-based de novo drug design using 3D deep generative models. Chemical Science, 12(41):13664–13675, 2021.

[42] Y. Li, L. Zhang, and Z. Liu. Multi-objective de novo drug design with conditional graph generative model. Journal of Cheminformatics, 10(1):33, July 2018.

[43] E. Lin, C.-H. Lin, and H.-Y. Lane. Relevant Applications of Generative Adversarial Networks in Drug Design and Discovery: Molecular De Novo Design, Dimensionality Reduction, and De Novo Peptide and Protein Design. Molecules, 25(14):3250, Jan. 2020.

[44] C. A. Lipinski, F. Lombardo, B. W. Dominy, and P. J. Feeney. Experimental and computational approaches to estimate solubility and permeability in drug discovery and development settings. Advanced Drug Delivery Reviews, 46(1-3):3–26, Mar. 2001.

[45] X. Liu, K. Ye, H. W. T. van Vlijmen, M. T. M. Emmerich, A. P. IJzerman, and G. J. P. van Westen. DrugEx v2: De novo design of drug molecules by Pareto-based multi-objective reinforcement learning in polypharmacology. Journal of Cheminformatics, 13(1):85, Nov. 2021.

[46] X. Liu, K. Ye, H. W. T. van Vlijmen, A. P. IJzerman, and G. J. P. van Westen. DrugEx v3: Scaffold-constrained drug design with graph transformer-based reinforcement learning. Journal of Cheminformatics, 15(1):24, Feb. 2023.

[47] Q. Long, C. Wu, X. Wang, L. Jiang, and J. Li. A Multiobjective Genetic Algorithm Based on a Discrete Selection Procedure. Mathematical Problems in Engineering, 2015:e349781, Sept. 2015.

[48] C. Lu, S. Liu, W. Shi, J. Yu, Z. Zhou, X. Zhang, X. Lu, F. Cai, N. Xia, and Y. Wang. Systemic evolutionary chemical space exploration for drug discovery. Journal of Cheminformatics, 14:19, Apr. 2022.

[49] J. Lyu, S. Wang, T. E. Balius, I. Singh, A. Levit, Y. S. Moroz, M. J. O’Meara, T. Che, E. Algaa, K. Tolmachova, A. A. Tolmachev, B. K. Shoichet, B. L. Roth, and J. J. Irwin. Ultra-large library docking for discovering new chemotypes. Nature, 566(7743):224–229, Feb. 2019.

[50] R. Macarron, M. N. Banks, D. Bojanic, D. J. Burns, D. A. Cirovic, T. Garyantes, D. V. S. Green, R. P. Hertzberg, W. P. Janzen, J. W. Paslay, U. Schopfer, and G. S. Sittampalam. Impact of high-throughput screening in biomedical research. Nature Reviews Drug Discovery, 10(3):188–195, Mar. 2011.

[51] A. Mauri and M. Bertola. AlvaBuilder: A Software for De Novo Molecular Design. Journal of Chemical Information and Modeling, July 2023.

[52] L. M. Mayr and D. Bojanic. Novel trends in high-throughput screening. Current Opinion in Pharmacology, 9(5):580–588, Oct. 2009.

[53] M. Mukaidaisi, A. Vu, K. Grantham, A. Tchagang, and Y. Li. Multi-Objective Drug Design Based on Graph-Fragment Molecular Representation and Deep Evolutionary Learning. Frontiers in Pharmacology, 13, 2022.

[54] A. Nigam, P. Friederich, M. Krenn, and A. Aspuru-Guzik. Augmenting Genetic Algorithms with Deep Neural Networks for Exploring the Chemical Space, Jan. 2020.

[55] A. Nigam, R. Pollice, M. Krenn, G. d. P. Gomes, and A. Aspuru-Guzik. Beyond generative models: Superfast traversal, optimization, novelty, exploration and discovery (STONED) algorithm for molecules using SELFIES. Chemical Science, 12(20):7079– 7090, May 2021.

[56] N. M. O’Boyle, M. Banck, C. A. James, C. Morley, T. Vandermeersch, and G. R. Hutchison. Open Babel: An open chemical toolbox. Journal of Cheminformatics, 3:33, Oct. 2011.

[57] M. Olivecrona, T. Blaschke, O. Engkvist, and H. Chen. Molecular de-novo design through deep reinforcement learning. Journal of Cheminformatics, 9(1):48, Sept. 2017.

[58] S.-s. Ou-Yang, J.-y. Lu, X.-q. Kong, Z.-j. Liang, C. Luo, and H. Jiang. Computational drug discovery Acta Pharmacologica Sinica, 33(9):1131–1140, Sept. 2012.

[59] P. Penner, V. Martiny, L. Bellmann, F. Flachsenberg, M. Gastreich, I. Theret, C. Meyer, and M. Rarey. FastGrow: On-the-fly growing and its application to DYRK1A. Journal of Computer-Aided Molecular Design, 36(9):639–651, Sept. 2022.

[60] D. A. Pereira and J. A. Williams. Origin and evolution of high throughput screening. British Journal of Pharmacology, 152(1):53–61, 2007.

[61] T. Pereira, M. Abbasi, B. Ribeiro, and J. P. Arrais. Diversity oriented Deep Reinforcement Learning for targeted molecule generation. Journal of Cheminformatics, 13(1):21, Mar. 2021.

[62] M. Podda, D. Bacciu, and A. Micheli. A Deep Generative Model for Fragment-Based Molecule Generation. In Proceedings of the Twenty Third International Conference on Artificial Intelligence and Statistics, pages 2240–2250. PMLR, June 2020.

[63] M. Popova, O. Isayev, and A. Tropsha. Deep reinforcement learning for de novo drug design. Science Advances, 4(7):eaap7885, July 2018.

[64] A. J. Price, S. Howard, and B. D. Cons. Fragment-based drug discovery and its application to challenging drug targets. Essays in Biochemistry, 61(5):475–484, Nov. 2017.

[65] D. C. Rees, Congreve, C. W. Murray, and R. Carr. Fragment-based lead discovery. Nature Reviews Drug Discovery, 3(8):660–672, Aug. 2004.

[66] S. Ryu and S. Lee. Accurate, reliable and interpretable solubility prediction of druglike molecules with attention pooling and Bayesian learning, Sept. 2022.

[67] A. V. Sadybekov and V. Katritch. Computational approaches streamlining drug discovery. Nature, 616(7958):673–685, Apr. 2023.

[68] M. F. Sanner. Python: A programming language for software integration and development. Journal of Molecular Graphics & Modelling, 17(1):57–61, Feb. 1999.

[69] B. Shaker, S. Ahmad, J. Lee, C. Jung, and D. Na. In silico methods and tools for drug discovery. Computers in Biology and Medicine, 137:104851, Oct. 2021.

[70] M. D. Shultz. Two Decades under the Influence of the Rule of Five and the Changing Properties of Approved Oral Drugs. Journal of Medicinal Chemistry, 62(4):1701–1714, Feb. 2019.

[71] M. Šícho, S. Luukkonen, H. W. van den Maagdenberg, L. Schoenmaker, O. J. M. Béquignon, and G. J. P. van Westen. DrugEx: Deep Learning Models and Tools for Exploration of Drug-Like Chemical Space. Journal of Chemical Information and Modeling, 63(12):3629–3636, June 2023.

[72] G. Sliwoski, S. Kothiwale, J. Meiler, and E. W. Lowe. Computational Methods in Drug Discovery. Pharmacological Reviews, 66(1):334–395, Jan. 2014.

[73] J. O. Spiegel and J. D. Durrant. AutoGrow4: An open-source genetic algorithm for de novo drug design and lead optimization. Journal of Cheminformatics, 12(1):25, Apr. 2020.

[74] N. Ståhl, G. Falkman, A. Karlsson, G. Mathiason, and J. Boström. Deep Reinforcement Learning for Multiparameter Optimization in de novo Drug Design. Journal of Chemical Information and Modeling, 59(7):3166–3176, July 2019.

[75] B. Tang, F. He, D. Liu, F. He, T. Wu, M. Fang, Z. Niu, Z. Wu, and D. Xu. AI-Aided Design of Novel Targeted Covalent Inhibitors against SARS-CoV-2. Biomolecules, 12(6):746, May 2022.

[76] G. W. Tang and R. B. Altman. Knowledge-based Fragment Binding Prediction. PLoS Computational Biology, 10(4):e1003589, Apr. 2014.

[77] T. F. Vieira and S. F. Sousa. Comparing AutoDock and Vina in Ligand/Decoy Discrimination for Virtual Screening. Applied Sciences, 9(21):4538, Jan. 2019.

[78] A. C. Walls, Y.-J. Park, M. A. Tortorici, A. Wall, A. T. McGuire, and D. Veesler. Structure, Function, and Antigenicity of the SARS-CoV-2 Spike Glycoprotein. Cell, 181(2):281–292.e6, Apr. 2020.

[79] D. Weininger. SMILES, a chemical language and information system. 1. Introduction to methodology and encoding rules. Journal of Chemical Information and Computer Sciences, 28(1):31–36, Feb. 1988.

[80] M. J. Wildey, A. Haunso, M. Tudor, M. Webb, and J. H. Connick. Chapter Five-High-Throughput Screening. In R. A. Goodnow, editor, Annual Reports in Medicinal Chemistry, volume 50 of Platform Technologies in Drug Discovery and Validation, pages 149–195. Academic Press, Jan. 2017.

[81] J. Wilson, B. A. Sokhansanj, W. C. Chong, R. Chandraghatgi, G. L. Rosen, and H.-F. Ji. Fragment databases from screened ligands for drug discovery (FDSL-DD). Journal of Molecular Graphics and Modelling, page 108669, Nov. 2023.

[82] D. Wolpert and W. Macready. No free lunch theorems for optimization. IEEE Transactions on Evolutionary Computation, 1(1):67–82, Apr. 1997.

[83] A. Yoshimori, F. Miljković, and J. Bajorath. Approach for the Design of Covalent Protein Kinase Inhibitors via Focused Deep Generative Modeling. Molecules (Basel, Switzerland), 27(2):570, Jan. 2022.

[84] T. Yoshizawa, S. Ishida, T. Sato, M. Ohta, T. Honma, and K. Terayama. Selective Inhibitor Design for Kinase Homologs Using Multi-objective Monte Carlo Tree Search. Journal of Chemical Information and Modeling, 62(22):5351–5360, Nov. 2022.

[85] W. Yu and A. D. MacKerell. Computer-Aided Drug Design Methods. Methods in molecular biology (Clifton, N.J.), 1520:85–106, 2017.

[86] Z. Zhou, S. Kearnes, L. Li, R. N. Zare, and P. Riley. Optimization of Molecules via Deep Reinforcement Learning. Scientific Reports, 9:10752, July 2019.

